# 3D Contractile and Remodeling Behaviors of Functionally Normal and Prolapsed Human Mitral Valve Interstitial Cells

**DOI:** 10.1101/2025.10.20.683556

**Authors:** Toni M. West, Gabriel Peery, Sanjana S. Chemuturi, Jodie H. Pham, Giovanni Ferrari, Michael S. Sacks

## Abstract

Mitral valve prolapse (MVP) can lead to heart failure, arrhythmia, and death. The only treatments available for MVP are replacement or repair; alternative therapies remain elusive due to lack of knowledge of the underlying pathological processes. The goal of the present study was thus to explore how MVP affects human mitral valve interstitial cell (hMVICs) extracellular matrix (ECM) remodeling and basal contractility characteristics. Isolated MVP and physiologically normal hMVICs were embedded in poly(ethylene) glycol-based hydrogels containing fluorescent fiducial markers and 3D traction force microscopy via inverse modeling was employed to determine the local change in hMVIC hydrogels due to enzymatic degradation and collagen deposition. Results indicated pronounced hydrogel softening occurred generally further from the hMVICs, whereas stiffening occurred in close proximity to hMVICs due to collagen deposition as verified by collagen-staining. MVP hMVICs induced greater hydrogel stiffening and less degradation than normal hMVICs. Interestingly, even though MVP hMVICs had higher basal contractile displacements, their corresponding traction forces and hydrogel strain energy densities were significantly lower than those of normal hMVICs. These findings elucidate, for the first time, that MVP hMVICs have significantly altered biophysical contractile and ECM remodeling behaviors compared to normal hMVICs.

**Simple Summary:** When a mitral heart valve gets thick, stiffened, and degraded, it can flip backwards (prolapse), causing blood to flow the wrong way. This can cause heart failure, arrhythmia, or even death. The only treatment for mitral valve prolapse (MVP) is surgery. To pave the way for a medication, this study aimed to understand how cells that maintain the mitral valve, mitral valve interstitial cells (MVICs), act on their surrounding tissue. The mechanical properties of the MVICs were tested, including how much they contract and how much they pull on their surroundings. Also, the mechanical properties that the MVICs place on their surroundings were tested, like how much they stiffen and degrade the tissue and how much energy is stored in the tissue as the MVICs contract. Compared to normal MVICs, we found that MVP MVICs stiffen their surroundings more and degrade their surroundings less. We also found that even though MVP MVICs contract more than normal MVICs, the energy that they place on their surroundings is less than that of normal MVICs, indicating that MVP MVICs are less mechanically effective. This is the first time that this ineffectiveness has been seen and may be key to targeting MVP with future medications.

## 1. Introduction

There are approximately 40,000 deaths worldwide annually from non-rheumatic primary-mitral valve disease, the most common type of which is mitral valve prolapse (MVP) [1]. In MVP, mitral valve (MV) interstitial cells (MVICs) not only increase their intracellular expression of *α*-smooth muscle actin (*α*SMA), but they also modulate their remodeling of the MV extracellular matrix (ECM), leading to tissue thickening and stiffening, excess collagen deposition and fragmentation, and increased matrix mealloproteinase (MMP) activity [2]. Once MVP in a patient has progressed to the point of having moderate to severe mitral regurgitation (MR) with concomitant symptoms (e.g., exercise intolerance or arrhythmia), surgical repair or replacement with their associated limitations are the only clinical treatment options [3]. Currently, there are no non-surgical therapies for MVP.

MV leaflets are composed primarily of a type-I collagen-rich ECM, which is produced and maintained by embedded MVICs. The degree of ECM remodeling that has occurred during MVP determines the degree of dysfunction [2]. Therefore, to move towards developing non-surgical (e.g., pharmacological) treatments for MVP, a better understanding of how human MVICs (hMVICs) affect their ECM in health and disease is greatly needed. The goal of the current work was thus to determine how hMVICs from MVP patients differentially alter their surroundings mechanically when compared to hMVICs from functionally normal mitral valves of donors.

A popular method to quantify the biophysical responses of cells is 3D traction force microscopy (TFM). In this approach, cells are embedded in a translucent hydrogel along with fluorescent microspheres which act as fiducial markers. The embedded cells are then exposed to biochemical agents to alter their contractile state, and the resulting movements of markers are then tracked [4]. From the geometry of the cell and marker displacements, inverse finite element models are then used to estimate the cellular traction forces [5–7]. However, usually these methods assume homogeneous hydrogel behaviors, ignoring the effects of cell-induced hydrogel degradation and ECM deposition. To address these issues, we have developed a pipeline where mathematical modeling of these heterogeneous phenomena can be performed [8,9]. These methods allow for determination of regional hydrogel softening due to enzymatic degradation and pronounced stiffening due to ECM deposition.

In the present study we hypothesized that since MVP MVICs produce additional *α*SMA [2], they would have increased basal contractility and thus generate greater traction forces. Additionally, as MVP hMVICs are known to produce both more collagen and more MMPs [2], we expected that both stiffer and more compliant hydrogel regions near the MVP MVICs than normal hMVICs. Furthermore, we expected that hMVICs would produce larger traction forces near cellular protrusions since most actin networks are seen in these areas of stained VICs [10]. Our results presented herein elucidate a more nuanced story.

## 2. Materials and Methods

hMVICs from mitral valve prolapse (MVP) patients and anatomically normal donors (Norm) that were embedded in crosslinked poly(ethylene) glycol (PEG)-based hydrogels containing fluorescent-microsphere fiducial markers were imaged in their basal and fully-relaxed states. The fiducial-marker displacements were then fit to the entire imaged volume by Gaussian process regression (GPR) to determine both basal hMVIC contractility and hydrogel displacement. Furthermore, the hydrogel displacements served as inputs for a detailed finite element inverse modeling approach that determined localized modulus changes resulting from enzymatic degradation and collagen deposition. The results of these computations also allowed for accurate calculation of hMVIC traction forces, as we did not need to assume the hydrogel had constant homogeneous mechanical properties.

### 2.1. hMVIC Sample Preparation and 3D-TFM Imaging

hMVICs were isolated as previously described [11,12] from three patients undergoing first-time surgery for mitral valve prolapse (MVP) and three anatomically normal (Norm) tissue donors. In brief, the endothelial cell layer was scraped from the leaflets, and the remaining tissue was minced and placed in 1 mg/ml collagenase type 2 solution (Worthington). Cells were then pelleted and resuspended in DMEM + 10% FBS + 1x PSG and grown on plastic to p1 to select for interstitial cells and frozen. Expansion was undergone on collagen-coated dishes to prevent non-phenotypic activation of the hMVICs [13]. p4 - p5 hMVICs were implemented for hydrogel embedding and imaging, which has been previously described [8,9]. To summarize, hMVICs stained with CellBrite Red were placed in a hydrogel precursor mix containing Dragon-green fluorescent microspheres (Bangs laboratories), MMP-degradable crosslinker (Bachem), CRGDS adhesion peptide (Bachem), and norbornene-functionalized 8-arm poly(ethylene) glycol (JenKemUSA) and were gelated with 3 minutes of UV. All materials were obtained from Thermo-Fisher unless otherwise stated. Hydrogels were incubated in DMEM + 10% FBS + 1x PSG for three days prior to imaging. A Nikon AXR laser scanning confocal microscope collected 3D images of hMVICs and the surrounding hydrogel fiducial markers in the basal and fully-relaxed, cytochalasin-D (CytoD) treated steady states as previously described [9,14]. The resulting images had isotropic voxels with side lengths of 0.288 *μ*m and overall volumetric dimensions of 147 × 147 × 119.23 *μ*m after spherical-aberration correction.

### 2.2. Immunocytochemistry Preparation and Imaging

hMVICs were encased in PEG hydrogel as described above but without Dragon green fiducial markers and fixed with 4% paraformaldehyde in dPBS after three days of culture. A solution of 15 *μ*g/mL wheat germ agglutinin (WGA) conjugated to Alexafluor-555 and 1:100 rabbit anti-collagen Type 1 antibody (Rockland 600-401-103) in Hank’s buffered salt solution (HBSS) was placed on the hydrogel overnight at 4°C for primary staining. Secondary staining occurred at 4°C overnight with a solution of 1:1,000 goat antirabbit alexafluor-488 in HBSS. Imaging occurred with the same microscope and settings as outlined above.

### 2.3. Displacement and Volumetric Analysis

Our custom developed software FM-TRACK [4] was implemented to track the displacements of the fiducial markers in the PEG hydrogel. As we have previously described [9], a Gaussian process regression (GPR) model of hydrogel displacements was fit for each cell system while accounting for the variance of noise in the input. Evaluation of the GPR model at the nodes of the hMVIC surface meshes was utilized to determine cellular surface displacements during basal contractility. Displacements oriented towards the center of the cell were identified and defined as basal contractile displacements. Additionally, a finite element mesh of first-order tetrahedral Lagrange elements was produced for the total imaged hydrogel volume of each system as previously described [9]. Nodal displacements in the meshes were determined with the GPR model. The resulting hydrogel displacements were then leveraged to calculate the Jacobian *J* = *det*(**F**) throughout each hydrogel, where **F** is the local deformation gradient tensor. Next, the hydrogel displacements were employed as inputs for the inverse model of heterogeneous hydrogel modulus.

### 2.4. Inverse Modeling of Heterogeneous Hydrogel Moduli

As we previously described [9], the hydrogels here were modeled as nonlinear materials that are neo-Hookean hyperelastic compressible solids, using the following spatially-varying strain energy density function:

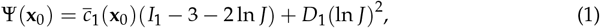

where **x**_0_ is the material point location in the hydrogel, *I*_1_ = tr **C** is the first invariant of the right Cauchy-Green tensor **C** = **F**^*T*^ **F**. The heterogeneous material-parameter field 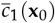 was further given an exponential form

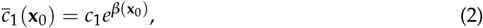

so that large orders of magnitude in modulus could be accounted for with adequate numerical stability. *c*_1_ was set to one half the mean (linear) shear modulus as measured by rheometry, 108 Pa/2 = 54 Pa. Based on previous results [9], *D*_1_/*c*_1_ = 1 was utilized to match the observed compressibility of the hydrogel. *β*(**x**_0_) was a field discretized in the first-order Lagrange basis of the hydrogel mesh whose degrees of freedom were control variables for optimization, with *β*(**x**_0_) = 0 as the initial guess, and forward solves were performed in Python FEniCS [15–22]. Inverse model optimization was performed with the limited-memory Broyden-Fletcher-Goldfarb-Shanno (L-BFGS) algorithm [23] to match displacements between target and simulation. The objective function that was minimized throughout this process with respect to *β*_*sim*_ was

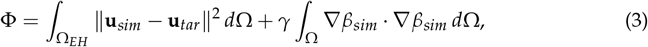

where **u**_*sim*_ and **u**_*tar*_ are the simulated and the target displacements, respectively. Ω is the imaged volume, and Ω_*EH*_ is the volume within the “event horizon” defined here as the volume in which fiducial-marker displacement magnitudes could be reliably determined relative to image voxel size, set to 0.38 *μ*m displacement magnitude cutoff as previously described [9]. Concentrating on Ω_*EH*_ additionally made the problem size more tractable for higher-end personal computers (i9 processor with 125 GB allocated RAM). Gradients of the objective functional, required by L-BFGS, were computed in FEniCS-adjoint [24] and MOOLA [25]. Simulated displacements at each iteration of inverse optimization were calculated by minimizing total hydrogel strain energy,

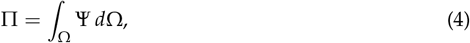

and Dirichlet boundary conditions were applied to the hMVIC surface and outer bounding box of the hydrogel, setting displacements there to observed values. The output of the full inverse model was the final modulus field prediction, where 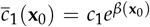, with convergence to these values being considered complete when either the objective function value fell below 10^−8^ or the magnitude of its gradient fell below 10^−6^. Error analysis of synthetic test cases determined that only 1.6% of nodes had errors ≥ 0.38 *μ*m, and runtime per hydrogel in a pilot study was approximately 1 hour [9].

### 2.5. Post-Processing

As has been previously outlined [9], the traction forces at the cellular surface were calculated by way of the first Piola-Kirchoff stress tensor, **P**. Distributions of strain energy density within the hydrogel during basal contractility were then determined and analyzed in an element-wise fashion. Graphs and related statistics were created with Python and matplotlib [26] and scipy [27] libraries. Visualizations were rendered in Paraview 5.13.2. p ≤ 0.05 was considered significant. 2-3 hMVICs/patient were analyzed, with each type of analysis employing the same hMVICs. Figure legends and text denote specific statistical tests applied in each graph or visualization.

## 3. Results

### 3.1. Basal Contractility Displacement

To determine basal contractility, a displacement field of the surrounding hydrogel of each hMVIC from the relaxed, Cyto-D treated state to the basal state was evaluated at the cellular surface (Figure 1a-b). The average basal contractile displacements of hMVICs from physiologically normal (Norm) and MVP mitral valves were compared (Table 1). The average effect size of 19.10% ((1.565 - 1.266)/1.565) and a p<0.001 depict that MVP hMVICs were under significantly larger basal contractile responses. This is in line with previous reports that MVICs have increased *α*SMA expression in MVP models and supports our hypothesis [28,29].

**Table 1.** Two-Tailed Unpaired T-tests with Welch’s Correction Results.

| Dependent Variable | Norm Mean ± SD | MVP Mean ± SD | *p* value |
| --- | --- | --- | --- |
| Contractile Displacement | 1.266 ± 1.077 μm | 1.565 ± 0.8742 μm | < 0.001 |
| Stiffened Moduli* | 374.6 ± 803.9 Pa | 445.7 ± 850.7 Pa | < 0.001 |
| Degraded Moduli* | 16.93 ± 8.9556 Pa | 19.66 ± 9.526 Pa | < 0.001 |
| Unsigned Surface Traction | 3.193 ± 14.94 kPa | 0.9351 ± 0.7008 kPa | < 0.001 |
| Event Horizon Strain Energy Density | 0.9552 ± 3.449 Pa | 0.7439 ± 2.339 Pa | < 0.001 |

\* Stiffened moduli are defined as those greater than median stiffening of 82 Pa and degraded moduli are those less than median degradation of 36 Pa.

**Figure 1.**
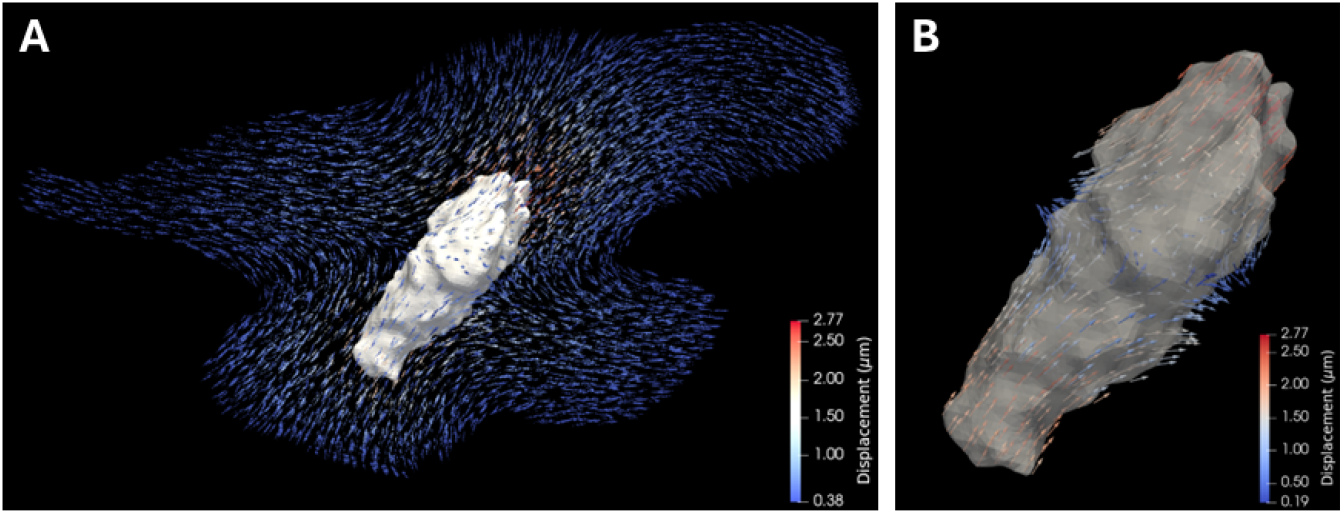
Representative hMVIC with basal contractility displacements. **a** Discretization of the displacement field at each hydrogel node within the event horizon of a representative hMVIC. **b** Cellular surface displacements of the same representative hMVIC.

### 3.2. Spatial Variation of J in ECM-Rich Hydrogels

Since we have seen previously that other hydrogel types were compressible [14], *J* was calculated on an element-wise basis in these PEG-based hydrogels and a comparison of variation in *J*(**x**_**0**_) was performed between the Norm and MVP groups (Figure 2). The PEG-based hydrogels portrayed large amounts of deformation, with 7.226% of mesh elements in all hydrogels having *J*(**x**_**0**_) ∉ [0.95, 1.05]. This verifies that PEG-based hydrogels undergo compressive behaviors and therefore must be treated as such in further calculations that will be delineated below. In addition, an Anderson-Darling test and an Earth mover’s distance test of the Norm vs. MVP groups for variation in *J*(**x**_**0**_) illustrated that there was a significant difference in *J*(**x**_**0**_) distribution between the groups, and it would take about a 1% change in all elements of a Norm hydrogel to become like an MVP hydrogel.

**Figure 2.**
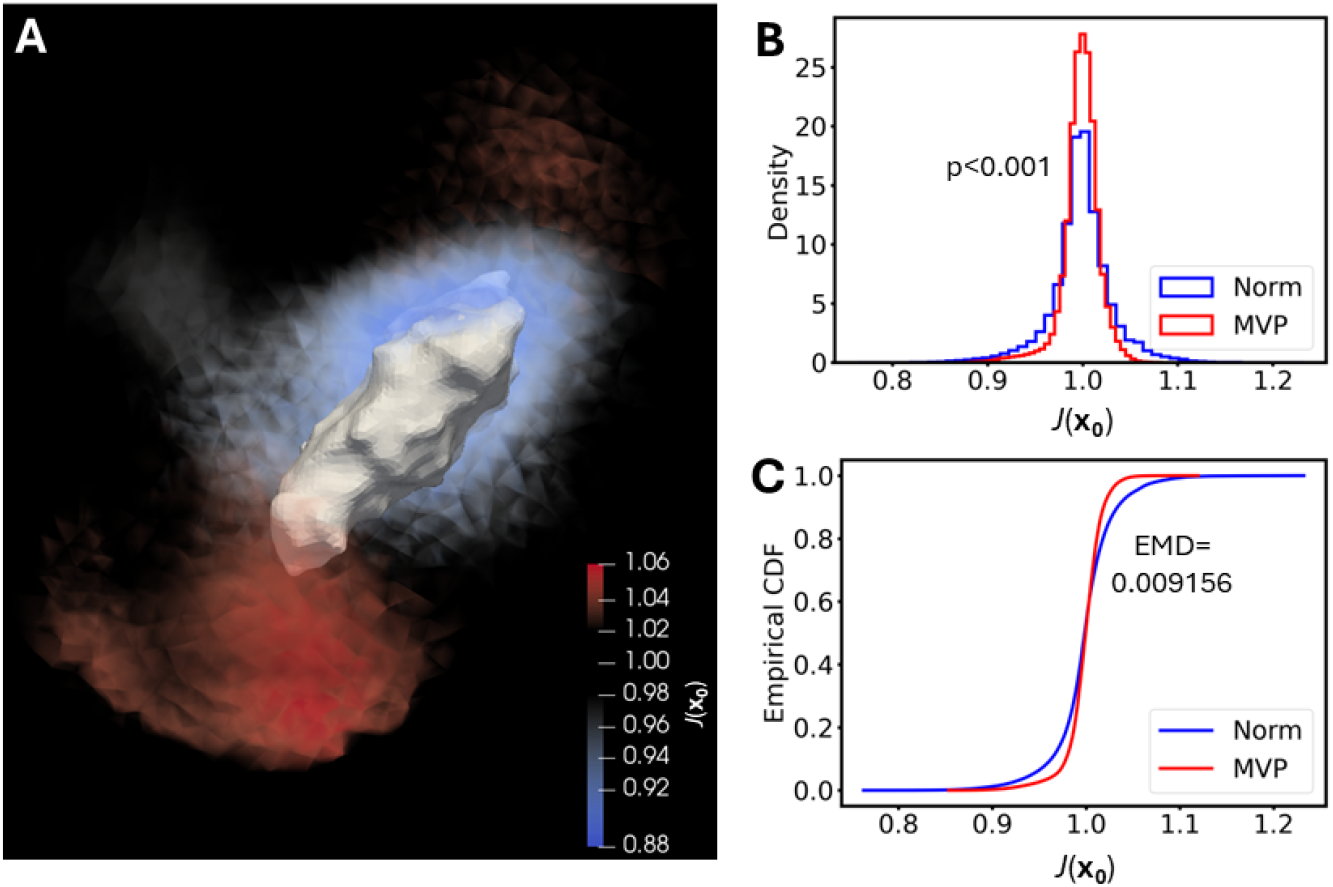
Large variations in *J*(x_0_) occur and are significantly different between anatomically normal and MVP groups. **a** Rendering of *J*(**x**_**0**_) within the hydrogel for a representative hMVIC. **b** Histogram and **c** empirical cumulative distribution function of *J*(**x**_**0**_) from the hydrogels surrounding all hMVICs per group. p<0.001 between groups by Anderson-Darling test. EMD=Earth mover’s distance test.

### 3.3. Cell-Mediated Extracellular Matrix Remodeling

To determine the extent to which the hMVICs had remodeled their surrounding hydrogels by excreting ECM components, the stiffness of the surroundings was modeled with an inverse method that determined heterogeneous neo-Hookean material parameters. The output moduli of the hMVIC-modified hydrogels were compared to collagen immunofluorescent staining of independent hMVICs (Figure 3). The elements of hydrogel that had greater than the global median of stiffening (83 Pa) correlated in pattern and approximate volume with what was seen in the collagen images, giving confidence that the inverse model modulus outputs capture biological processes of ECM modification.

**Figure 3.**
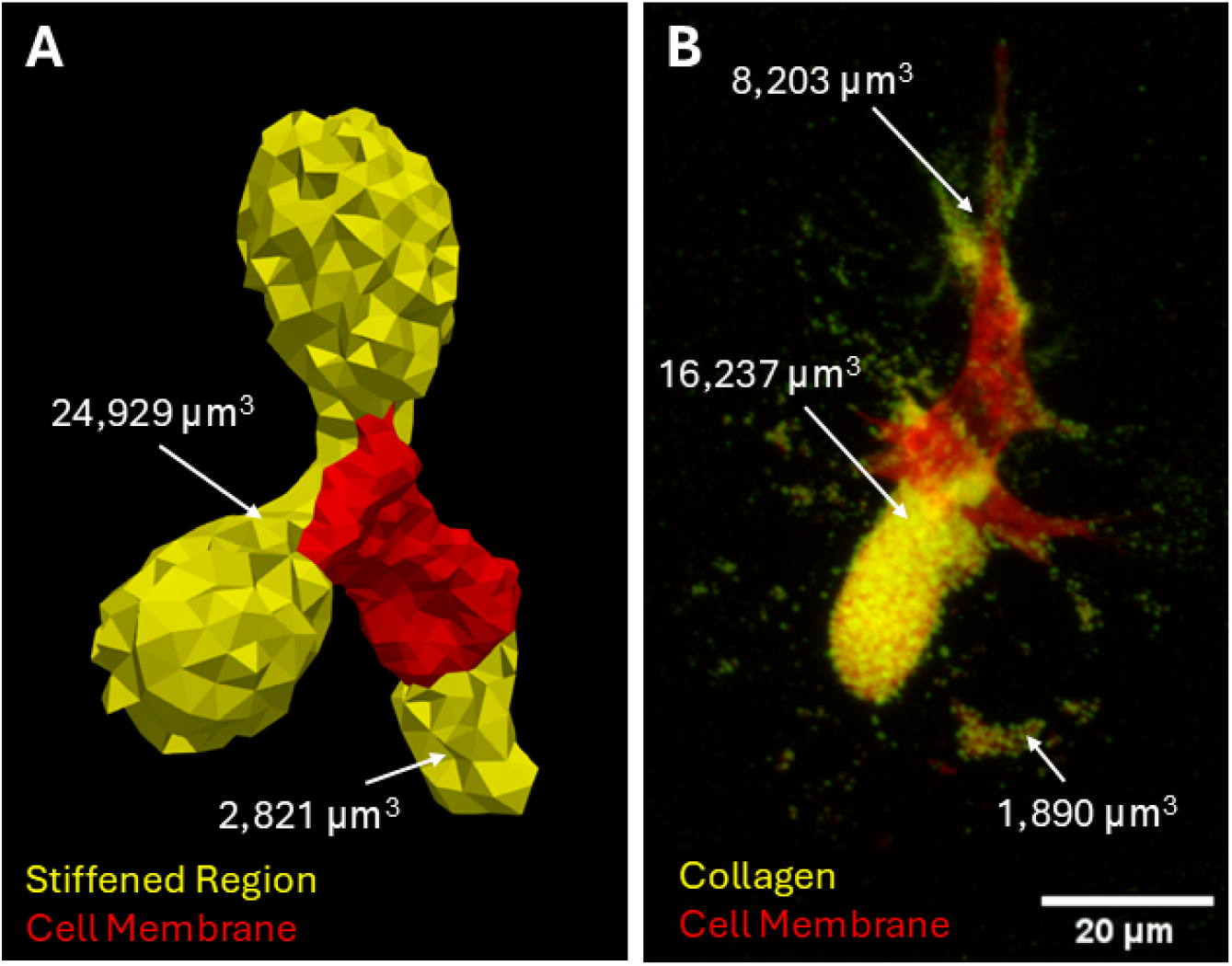
Stiffened hydrogel regions and stained collagen correlate in placement and volume. **a** Rendering of the zones of hydrogel around a representative hMVIC that have higher than the global median stiffened value of 82 Pa, as calculated by the inverse model. Total stiffened volume = 27,750 *μ*m^3^. **b** 2D standard deviation projection of a 3D image of an hMVIC stained for extracellular collagen. Volumes of individual zones of collagen surrounding the hMVIC were analyzed in full 3D in ImageJ-Fiji. Total collagen volume = 26,330 *μ*m^3^. Color legend and scale bar depicted in images.

The resulting moduli of each hydrogel model were subdivided into zones where greater than median stiffening had occurred (> 82 Pa) and degradation had occurred (< 36 Pa; Figure 4a). The hydrogel elements of stiffening and degradation were then separately compared between Norm and MVP groups. The stiffened regions near MVP hMVICs possessed significantly stiffer moduli than these regions near Norm hMVICs (Table 1). This supports our hypothesis and is in line with previous reports that state activated MVICs increase deposition of stiff ECM components such as collagen [2]. But surprisingly, the degraded regions of MVP hMVIC hydrogels had significantly less degradation than Norm hMVIC hydrogels (Table 1). In the degraded regions, the MVP average hydrogel modulus only fell 34.44 Pa over three days of modification (54 Pa unmodified hydrogel - 19.66 Pa average = 34.44 Pa) while the Norm hydrogel modulus fell 37.07 Pa. This data depicts that there is an additional 4.870% decrease in modulus in Norm hydrogels over three days of culture compared to that of MVP hydrogels. Therefore, MVP hMVICs produce stiffer ECM both due to an increase in stiffening and a decrease in degradation.

**Figure 4.**
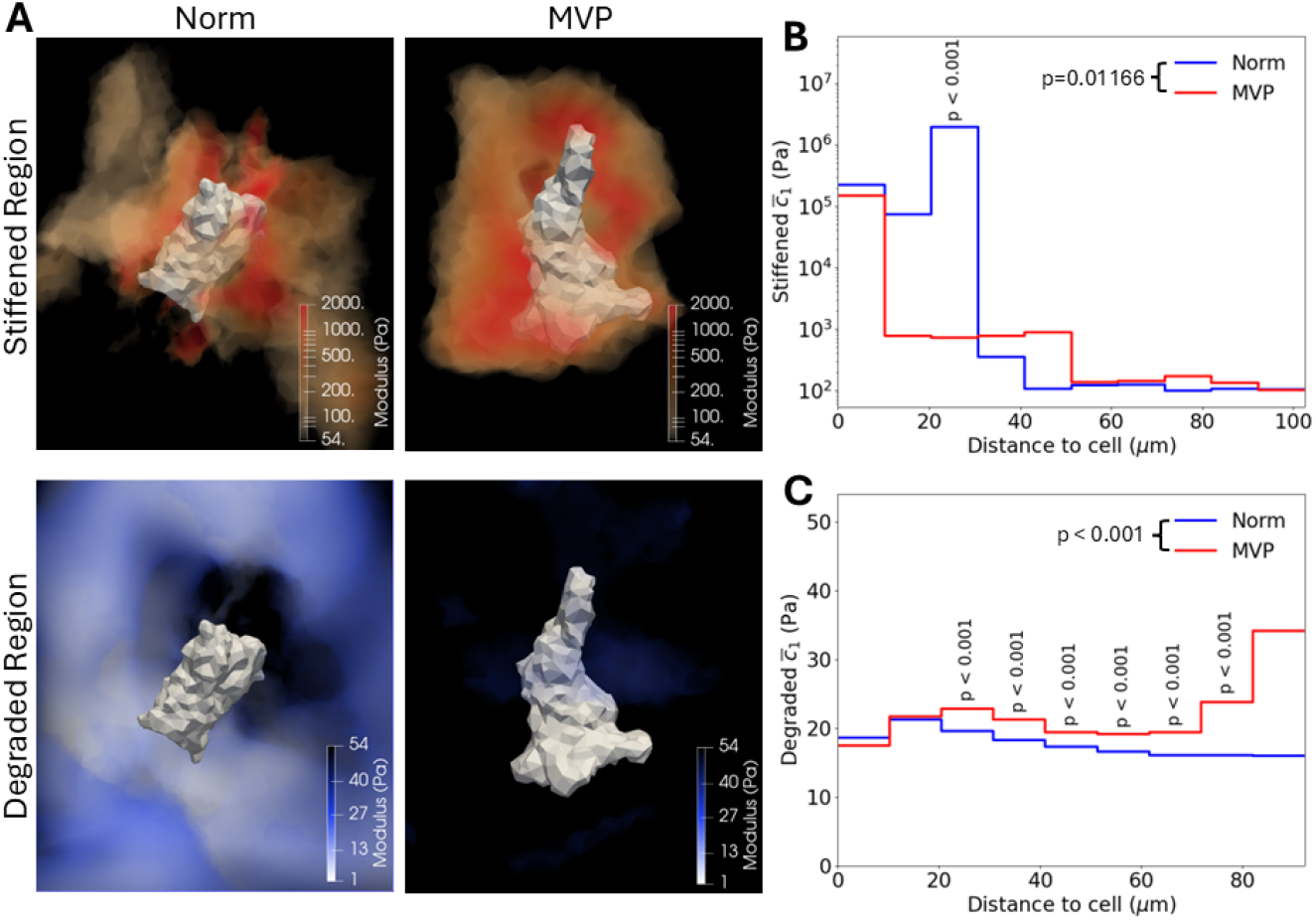
MVP hMVICs produce stiffer extracellular matrices. **a** Representative renderings of stiffened and degraded regions of hydrogels modified by hMVICs from physiologically normal (Norm) and mitral valve prolapse (MVP) valves. **b-c** Step graphs of stiffened moduli (>82 Pa; **b**) or degraded moduli (<36 Pa; **c**) with respect to distance bins of 10 *μ*m each. *p* values on legend calculated by two-way ANOVA. *p* values above steps calculated by Tukey post-test between Norm and MVP groups at that step.

Since visual inspection of the data suggested the stiffened areas were in close proximity to the hMVIC surfaces, an analysis was performed where the moduli were placed in 10 bins according to their distance away from the cell, with stiffened moduli and degraded moduli graphed separately. Stiffening trended towards being higher near the cell surface, but distance was not a significant variable for determining modulus (*p*=0.09257) by two-way ANOVA due to large variations in stiffened moduli (Figure 4b; Table S1). Still, significant differences in moduli could be seen between the differing distance bins. Normal hMVICs stiffened their surroundings approximately 2,000 times the unaltered hydrogel modulus within the first 20 *μ*m, which then significantly increased to approximately 20,000 times the unaltered modulus between 20 - 30 *μ*m (p < 0.001 by Tukey post-test). At 30-40 *μ*m away, hydrogels modified by normal hMVICs plummeted in stiffness to approximately 6 times the unmodified hydrogel modulus (*p*=0.005 by Tukey post-test) and then moved to approximately 2 times the unmodified modulus starting at 40 *μ*m. Alternatively, MVP hMVICs only stiffen to the 2,000x level for the first 10 *μ*m but then continue to stiffen at the more moderate 15x level for 10 - 50 *μ*m of hydrogel and then drop to about 2 times the unmodified modulus further away.

Degradation with respect to distance from the hMVIC was similarly assessed (Figure 4c; Table S2). The two-way ANOVA with Tukey post-test revealed that degradation stays steady in normal hMVICs with distance, causing approximately 40 Pa of change in stiffness from the unaltered hydrogel modulus, no matter the location. In contrast, MVP hMVICs have a significant decrease in the amount of degradation with increasing distance from the hMVIC, which is a significantly different pattern from the Norm hMVICs (*p* < 0.001 by two-way ANOVA). This highlights that the overall decrease in degradation potential that MVP hMVICs portrayed is mostly due to a decrease in degradation far away from the cell boundary.

### 3.4. Cellular Traction Forces

The traction forces were found to be significantly lower in MVP hMVICs when compared to normal hMVICs, suggesting a different mechanism than originally anticipated (Figure 5; Table 1). Indeed, when the traction-to-displacement ratio (TDR) was calculated, MVP hMVICs were approximately 76% less effective at transmitting traction to their surroundings (Norm: 3.193/1.266 = 2.522; MVP: 0.9351/1.565 = 0.5975). Renderings of each hMVIC were then reviewed to look for possible trends. When any hMVIC displaces during basal contraction, it tends to contract at the protrusions and at the same time have an isovolumetric expansion in the body of the cell (Figure 1b). Renderings of the hMVICs indicated that large traction forces often occurred near protrusions where contraction occurred, so a break down of the traction forces by contractile and expansile regions of the hMVICs was conducted (Figure 5; Table 2). In both contractile and expansile regions, MVP hMVICs had significantly lower average traction than normal hMVICs. Interestingly, there was a switch in the dominance of specific regions. In normal hMVICs, larger tractions occur in the expansile regions than in the contractile regions, while in MVP hMVICs, larger tractions occur in the contractile regions than the expansile regions, suggesting that cell contractile polarity may be altered during disease progression.

**Table 2.** MVP hMVICs exert significantly less unsigned traction in both contractile and expansile cellular regions.

| Traction |  |  |  |
| --- | --- | --- | --- |
|  | Norm Mean ± SD | MVP Mean ± SD | p value* |
| Contractile | 4.081 ± 27.02 kPa | 2.148 ± 20.19 kPa | 0.01450 |
| Expansile | 9.418 ± 44.01 kPa | 0.3360 ± 2.496 kPa | < 0.001 |
| p value* | <0.001 | 0.7948 |  |

\*p values calculated between corresponding row or column comparison by two-way ANOVA with Tukey post-test.

**Figure 5.**
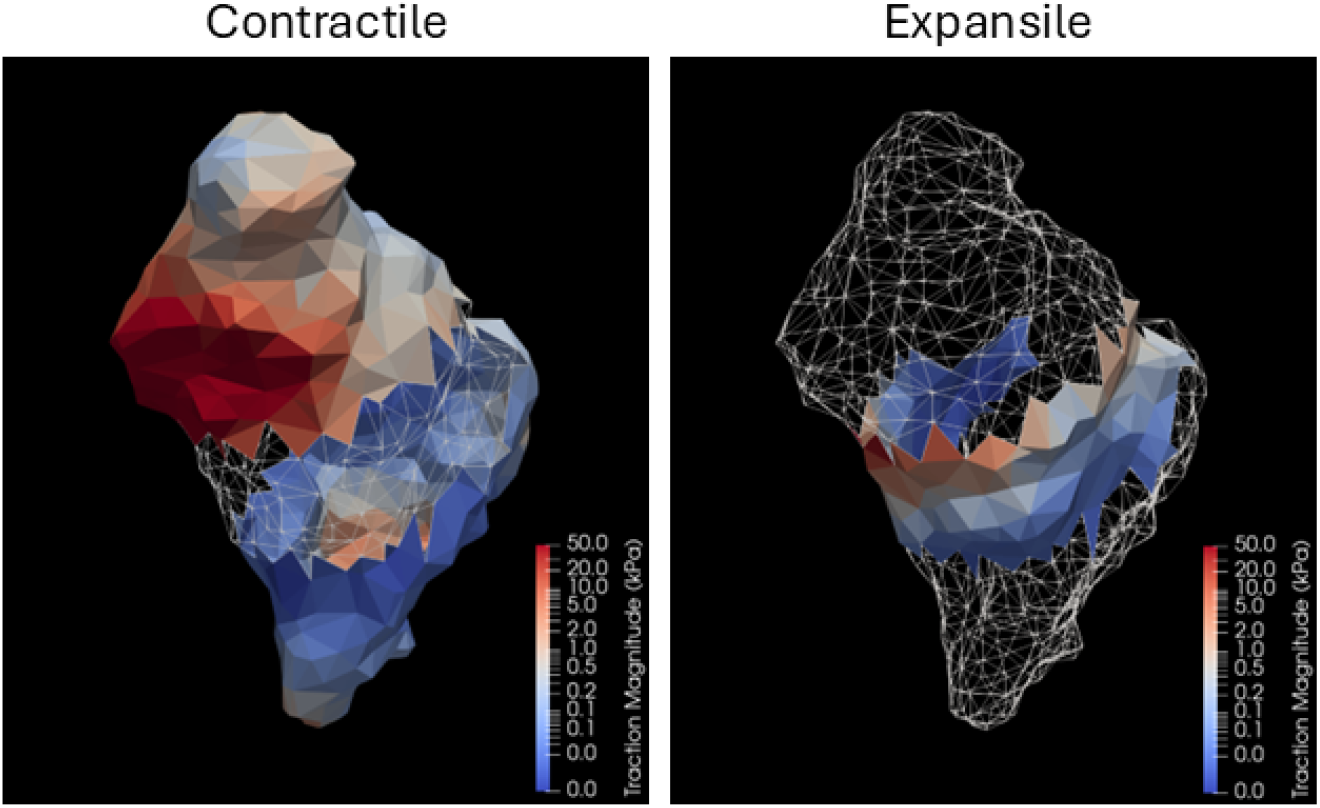
Traction forces in the contractile and expansile regions of a representative hMVIC. Rendering of traction forces present on each surface element of an example hMVIC depicting the elements where the cell was contracting inward towards the hMVIC centroid (Contractile) or expanding outward from the centroid (Expansile).

### 3.5. Strain Energy Density

To further investigate the effectiveness of how the traction forces from the hMVICs affect their surroundings, the strain energy density of the hMVIC-modified hydrogel was determined. Strain energy is a particularly attractive biophysical quantity that represents the total energy of cellular contraction, including contractile and gel mechanics. As expected, in all hMVICs, strain energy density was highest near the hMVIC boundary and significantly decreased with increasing distance from the cell (Figure 6). Similar to that seen with traction forces, the average of strain energy density from the MVP hMVIC ECM was lower than that of the normal hMVIC hydrogels (Figure 6; Table S3), which was mostly due to a decrease in strain energy density 10 - 30 *μ*m away from the cell membrane in MVP hMVIC ECM. But notably, MVP hMVICs produced a significantly higher strain energy density within the first 10 *μ*m from the cell boundary. Any difference that occurred between Norm and MVP hMVIC hydrogels in strain energy density further than 30 *μ*m away from the cell boundary were not significant and trended towards 0. Also as expected, stiffened areas of ECM had significantly higher strain energy density than degraded regions (Table 3). Since contractile polarity of cell membrane traction force was altered in disease, an investigation of whether this polarity shift was translated to strain energy density of the surroundings was then conducted. Both hydrogels modified by normal and MVP hMVICs exhibited significantly higher strain energy density in contractile regions than in expansile regions. Furthermore, stiffened MVP hMVIC ECM exhibited significantly lower strain energy than Norm hMVIC ECM, suggesting that the changes in overall strain energy density throughout the MVP ECM are mostly governed by changes that occur in the stiffened regions.

**Table 3.**
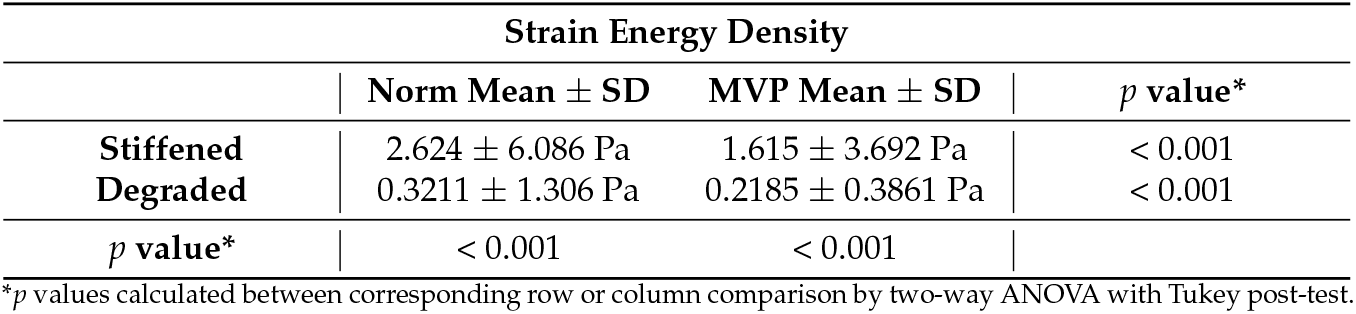
Strain energy density is higher in stiffened hydrogel zones and is lessened by MVP status.

**Table 4.**
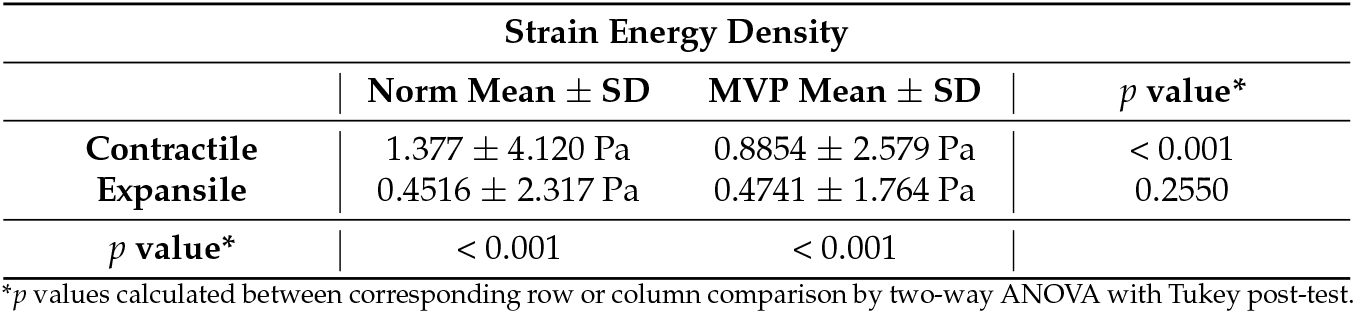
Strain energy density is higher in contractile hydrogel zones and is lessened by MVP status.

**Figure 6.**
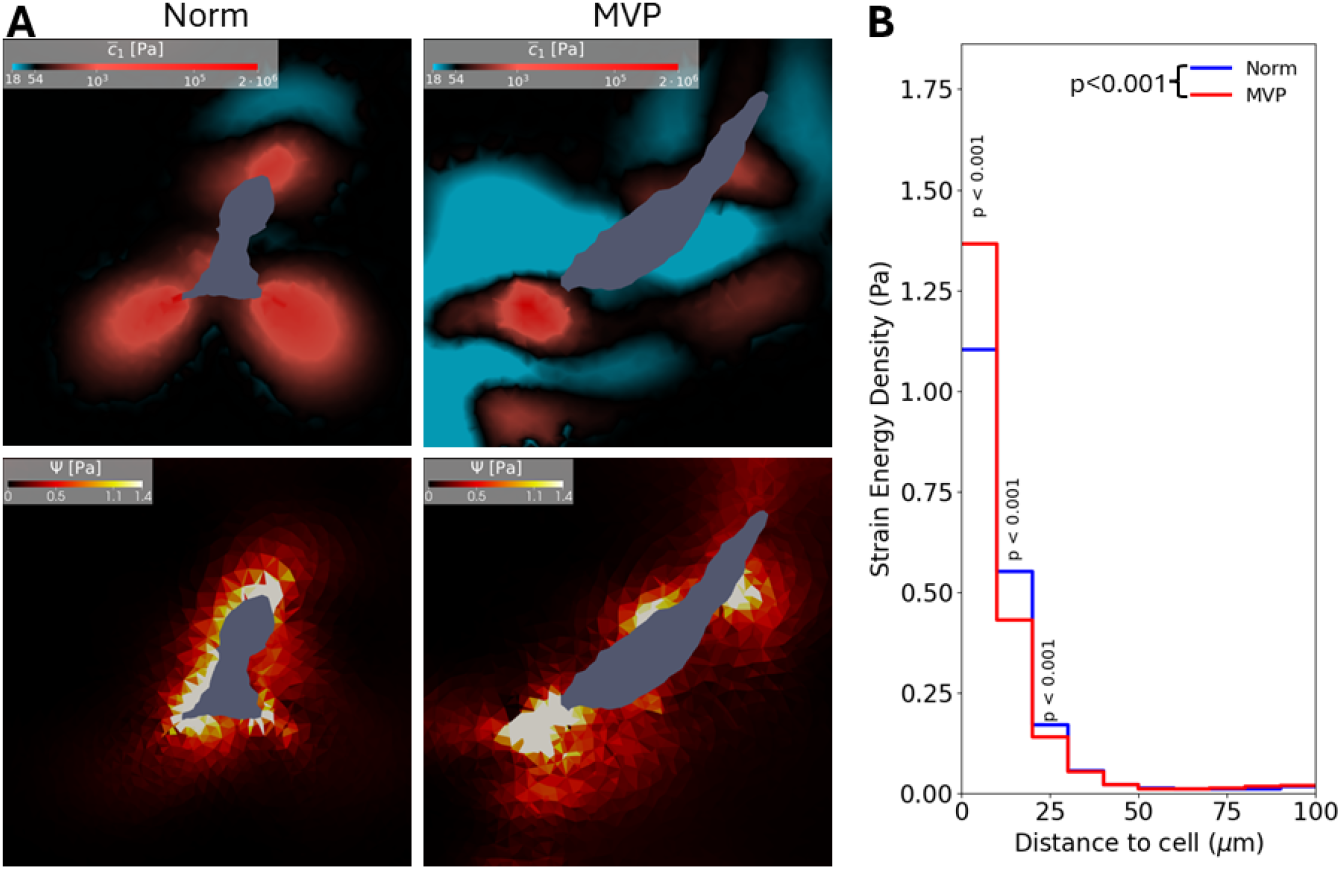
Strain energy density decreases with increased distance from the hMVIC and correlates with regions of high stiffness. **a** Cross-section slices of a normal (Norm) and MVP hMVIC where the stiffened moduli 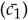 and high strain energy density (Ψ) align. **b** Step graph of strain energy density vs binned distances from the cell boundary. *p* value depicted in legend determined by two-way ANOVA. *p* values depicted above steps determined by Tukey post-test.

## 4. Discussion

In this investigation it was found that hMVICs from MVP patients produced significantly-stiffer and less-deformable matrices whose stiffening was more far-reaching from the cell. And although MVP hMVICs had significantly larger basal contractile displacements than Norm hMVICs, MVP hMVICs produced significantly less traction force at the cellular surface and less strain energy density within the hydrogels they modified.

One aspect of the ECM remodeling that was importantly different between the MVP and normal groups was that the MVP hMVIC hydrogels were less degraded, especially at greater distances from the cell boundary. Since the PEG hydrogel formulation used in this investigation is specifically designed to allow for MMP degradation [8], it was assumed that the difference between MVP and normal hydrogel degradation was due to a difference in hMVIC-mediated MMP regulation. Certain secretory MMPs are known to be up-regulated during MVP including MMPs 1, 2, 3, 7, and 9 [2], so it was hypothesized that MVP hMVIC hydrogels would have higher rates of degradation than hydrogels from normal hMVICs. The current data analysis did not support this hypothesis. A review of the literature revealed that because MMPs have relatively small molecular weights compared to other components of ECM or hydrogel matrices, MMPs can move through the matrix over time by Graham’s law of diffusion, but are also affected by the electrochemical and binding interaction between the MMP and the matrix substances [30,31]. In the MVP hMVIC hydrogels, there was a significantly lower amount of degradation further away from the cell. This rise in modulus with distance may be explained by a shift in expression to larger molecular weight secretory MMPs such as MMP-9 (92 kDa) or to MMPs with high affinity to native collagen, like MMP-1 [32,33]. Therefore, future fluorescence correlation spectroscopy experiments looking into the diffusion of MMPs from MVICs with MMP-1 or MMP-9 knockdown may elucidate how MMP expression profile in MVICs plays a role during MVP-related leaflet degradation [31].

The quintessential marker of MVP in MVICs is increased *α*SMA expression [2]. Because this is well substantiated, it was hypothesized that increases of *α*SMA in MVP hMVICs would lead to increased cell surface traction forces and hydrogel strain energy density. But this was not the case. Both basal traction and basal strain energy density were measured to be significantly lower in MVP samples. There are several possible mechanisms that could be occurring to lead to these unexpected results. Firstly, there is not a 1:1 ratio of *α*SMA expression to its incorporation into polymerized actin stress fibers. For instance, TGF*β* and ERK signaling up-regulate *α*SMA expression via SMAD and MAPK pathways while *α*SMA incorporation into polymerizing stress fibers is largely mediated by RhoA/ROCK signaling [2,34]. The latter is more influenced by mechanobiological inputs such as substrate stiffness and intracellular tension than it is by non-canonical TGF*β* and ERK signaling [34,35]. Therefore, it is possible that although there is increased expression of *α*SMA, there could be a lack of polymerization of it into actin stress fibers thereafter, which would explain how lower traction forces were seen in MVP hMVICs here.

The lack of *α*SMA polymerization may be in part due to the bidirectional feedback loop between focal adhesion stability and stress fiber formation described in fibroblasts, which occurs through ROCK signaling and likely translates to MVICs [35]. Despite being embedded in locally equal or stiffer hydrogel regions, MVP hMVICs generated lower cell surface traction forces and exerted less strain energy than normal hMVICs, indicating an uncoupling of their adhesion mechanisms. Further investigations into which of these mechanisms are important for differential MVP hMVIC mechanobiological phenotypes are therefore warranted. Several possibilities for uncoupling are likely, including reduced integrin expression [36], reduced integrin-ECM binding activity [37], reduced attachment of adhesions to the cytoskeleton *via* talin and vinculin [38], and reduction in organization of the deposited ECM, the intracellular cytoskeleton, or cellular morphology [39–41]. MVP hMVICs displayed a shift in traction force dominance from expansile to contractile regions relative to normal cells, highlighting a potential role for altered cell morphology in the disease phenotype. In addition to actin and focal adhesion components, other cytoskeletal elements such as microtubules may also influence traction generation and intracellular strain, motivating further investigation into their role in MVP hMVIC mechanobiology. These differences in cell-generated traction and polarity may ultimately influence how hMVICs remodel and stiffen their surrounding ECM, motivating closer examination of local strain energy distributions.

Interestingly, the strain energy densities of the closest areas to the cell boundaries were actually higher in hydrogels modified by normal hMVICs than in those modified by MVP hMVICs, which then drops off more quickly in normal than in MVP samples. This suggests that MVP hMVICs may have an impaired ability to mature the ECM macroproteins that they secrete. This aligns with what is seen in histology cross sections of mitral valve leaflets, where MVP leaflets have a more unstructured ECM than normal leaflets [2]. Our future endeavors therefore involve further developing the inverse modeling pipeline for modulus determination so that anisotropic ECM modification can be elucidated in the analysis.

### 4.1. Limitations

This study was able to determine the heterogeneous values of basal displacements, deformations, moduli, traction forces, and strain energy densities surrounding hMVICs from normal and MVP patients. Due to experimental noise and the need to make the problem size tractable, only hydrogel displacements greater than 0.38 *μ*m were reliably assessed. Although this represents very fine spatial resolution, some hMVICs had to be excluded because the assessable volume was too small. While this may have slightly affected the absolute average values, the observed trends between normal and MVP hMVICs remain valid, as the same 0.38 *μ*m cutoff was applied consistently across all analyses. When examining computed moduli, a liberal cutoff at the global median for stiffened or degraded regions was applied to minimize noise. Future work will aim to refine the cutoffs of 82 Pa and 36 Pa to values closer to the unmodified hydrogel modulus of 54 Pa. Nonetheless, the current methodology provides a rich quantitative understanding of ECM-rich hydrogel mechanobiology. Even small collagen deposits away from the cell body (likely laid down during cell migration [42–44]) were detectable implementing this workflow. Although there are inherent limitations, these models accurately capture key biological processes and reveal mechanical properties of hMVICs that cannot be directly measured. Importantly, this analysis demonstrates that MVP hMVICs exhibit a paradoxical phenotype: they produce a stiffer ECM while imparting lower traction forces and strain energy densities onto that ECM.

## 5. Conclusion

This study aimed to understand how hMVICs affect their ECM both through ECM remodeling and basal contractility. By conducting such a study, the differences in biomechanical mechanisms that change in hMVICs and their surrounding ECM during MVP were discerned. Through the implementation of 3D traction force microscopy [8,14] and our inverse modeling process [9], it was seen that MVP hMVICs have larger basal contractile displacement and remodel their surroundings to become significantly stiffer. At the same time, MVP hMVICs imparted significantly less traction force which led to significantly less strain energy density in the surrounding ECM-rich hydrogels of MVP hMVICs. These results are in line with what is seen on an organ level during MVP, where leaflet thickening and stiffening, and ECM disorganization are hallmarks of the disease [2]. Further investigation as to how MVP hMVICs concurrently impart less traction and strain energy than normal cells even though they have higher expression of actin polymers including *α*SMA [2] is clearly warranted. Understanding this dichotomy that is seen here for the first time may be an important link in elucidating how MVP hMVICs progress into their disease state and how we can mitigate this progression biochemically.

## Abbreviations

The following abbreviations are used in this manuscript:

3D-TFM: three-dimensional traction force microscopy
*α*SMA: alpha-smooth muscle actin
CytoD: cytochalasin-D
dPBS: Dulbecco’s phosphate buffered saline
ECM: extracellular matrix
MVP: mitral valve prolapse
ERK: extracellular signal regulated kinase
GPR: Gaussian process regression
HBSS: Hank’s buffered salt solution
hMVICs: human mitral valve interstitial cells
MAPK: mitogen-activated protein kinase
MMP: matrix metalloproteinase
MR: mitral regurgitation
MV: mitral valve
MVICs: mitral valve interstitial cells
MVP: mitral valve prolapse
Norm: physiologically normal
PEG: poly(ethylene) glycol
RhoA: ras homolog gene family, member A, GTPase
ROCK: rho-associated coiled-coil kinase
SMAD: SMA/MAD homologous protein
TDR: traction-to-displacement ratio
TGF*β*: transforming growth factor *β*
WGA: wheat germ agglutinin

## Author Contributions

Conceptualization: MSS (lead) and TMW (supporting), Experimental methodology: TMW (lead), Analytical and overall computational approach: MSS (lead) and GP (supporting); Software Development: GP (lead) and TMW (supporting); validation: GP (equal) and TMW (equal); formal analysis: TMW (equal), GP (equal), SSC (supporting), and JHP (supporting); investigation: TMW (equal), GP (equal), SSC (supporting), and JHP (supporting); resources: GF (lead) and MSS (supporting); data curation: TMW (equal), GF (equal), GP (supporting), SSC (supporting), and JHP (supporting); writing—original draft preparation: TMW (lead); writing—review and editing: MSS (lead), GF (supporting), GP (supporting), SSC (supporting), and JHP (supporting); visualization: TMW (equal) and GP (equal); supervision: MSS (lead) and TMW (supporting); project administration: MSS (lead), GF (supporting) and TMW (supporting); funding acquisition: MSS (equal), GF (equal), TMW (equal), and GP (equal). All authors have read and agreed to the published version of the manuscript.

## Institutional Review Board Statement

All human research conducted within this study conformed to the principles of the Declaration of Helsinki and were approved by Columbia University (IRB AC-AABC3510) and the University of Texas at Austin (IBC-2023-00293). All patient information was deidentified.

## Informed Consent Statement

Patients with mitral valve prolapse that were referred for first-time surgery were enrolled in the study with written informed consent at Columbia University Irving Medical Center and their MV leaflets were collected during a standard-of-care procedure. Anatomically normal MV leaflets were collected from hearts obtained from the heart transplant service of the Hospital of the University of Pennsylvania with IRB approval (IRB 802781) that were unable to be implanted at the time of organ donation and the donor had consented to donate tissue for research purposes in writing on Live On New York.

## Data Availability Statement

The raw data supporting the conclusions of this article will be made available by the authors on request.

## Acknowledgments

All microscopy was performed at the Center for Biomedical Research Support Microscopy and Flow Cytometry Facility at the University of Texas at Austin (RRID: *SCR*021756). During the preparation of this manuscript/study, the author(s) minimally utilized ChatGPT-5o for the purposes of manuscript word choice and code debugging. The authors have reviewed and edited the output and take full responsibility for the content of this publication.

## Supplementary Materials

**Table S1.**
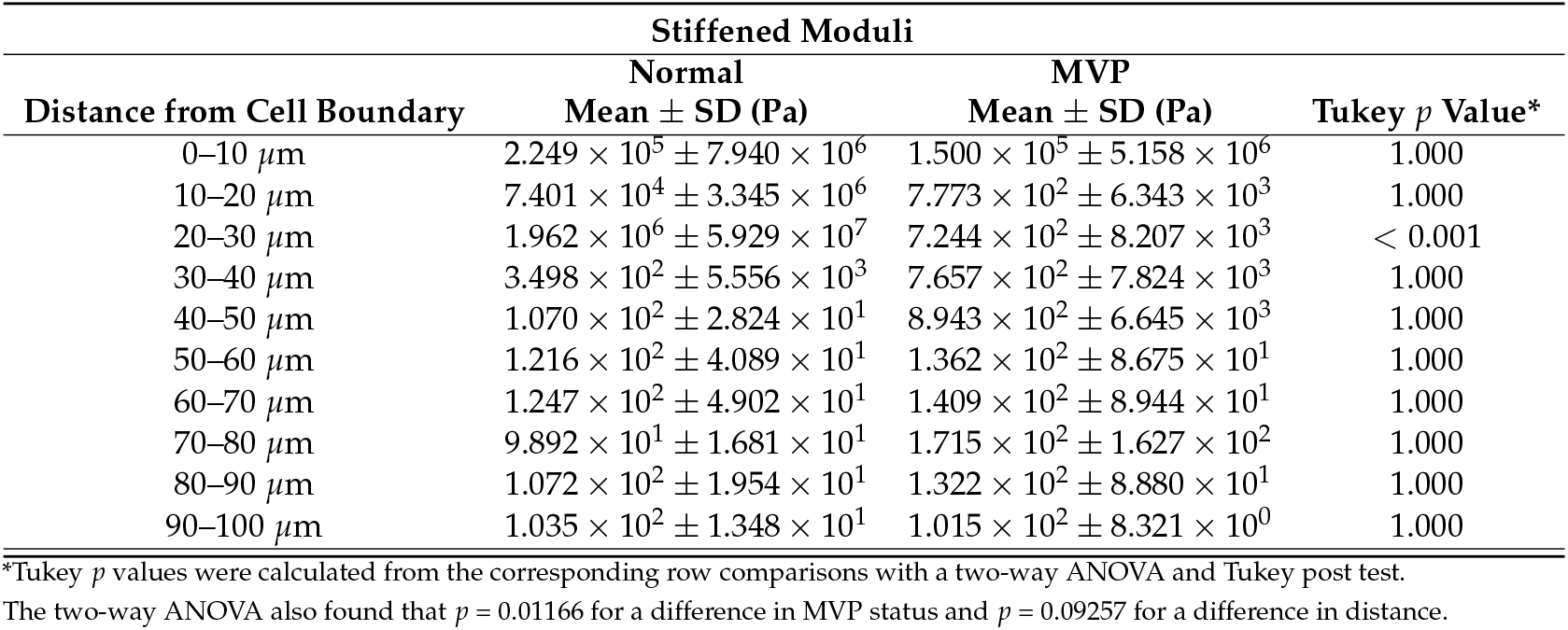
Stiffened Moduli by Distance Significantly Change by MVP Status.

**Table S2.**
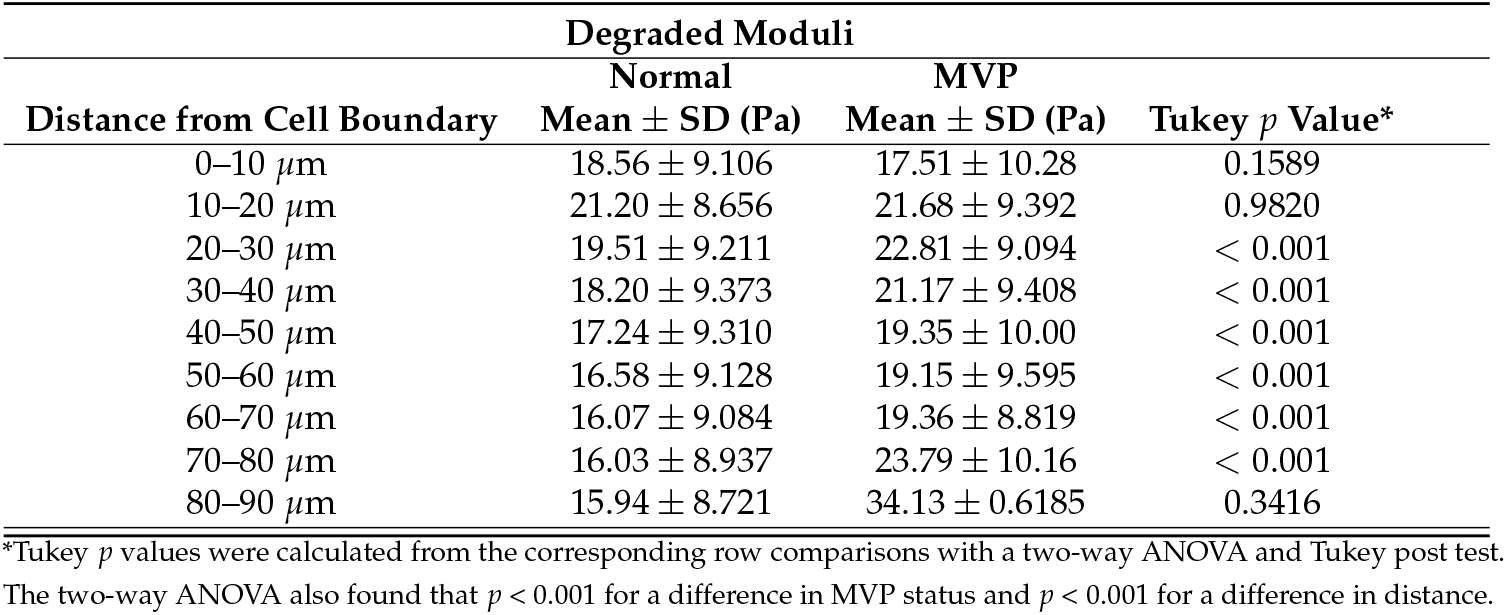
Degraded Moduli Significantly Changes by MVP Status and Distance from the Cell Boundary.

**Table S3.**
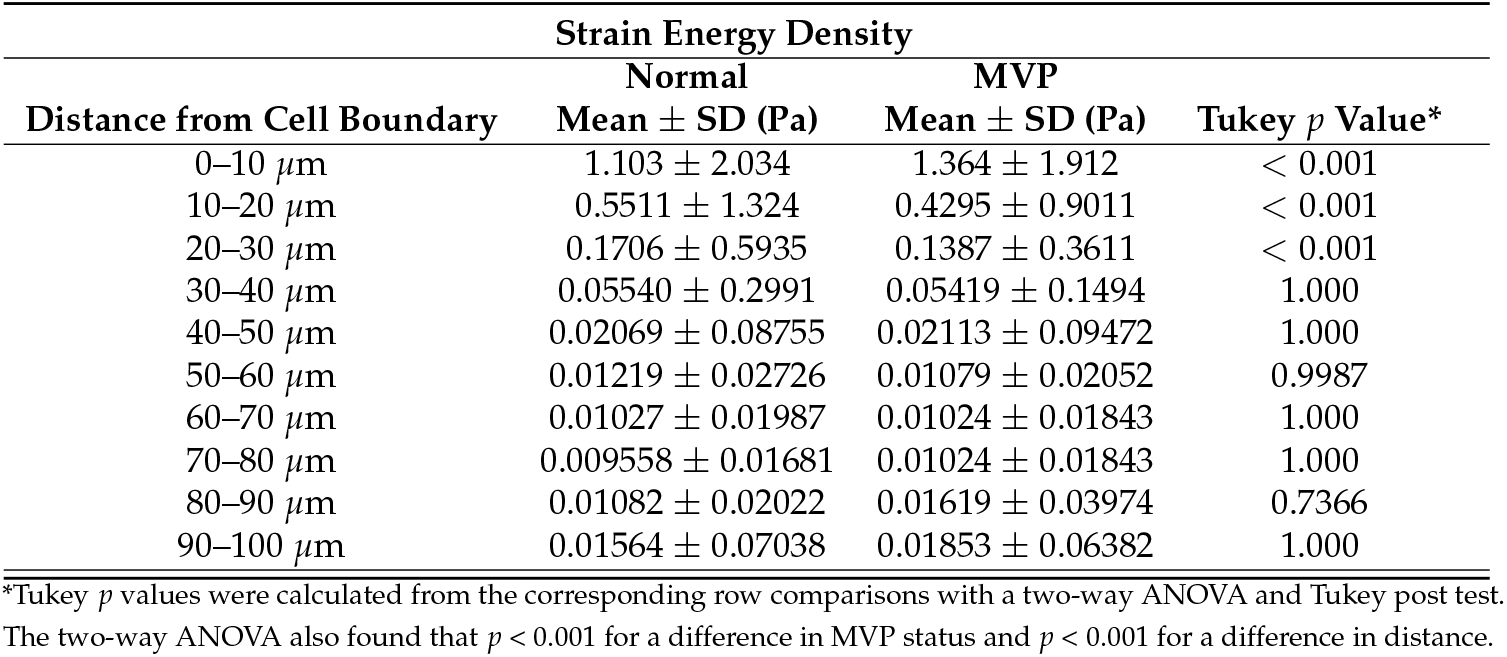
Hydrogel Strain Energy Density Significantly Changes by MVP Status and Distance from the Cell Boundary.

## Disclaimer/Publisher’s Note

The statements, opinions and data contained in all publications are solely those of the individual author(s) and contributor(s) and not of MDPI and/or the editor(s). MDPI and/or the editor(s) disclaim responsibility for any injury to people or property resulting from any ideas, methods, instructions or products referred to in the content.

