## Supplemental Tables for "3D Contractile and Remodeling Behaviors of Functionally Normal and Prolapsed Human Mitral Valve Interstitial Cells"

**Table S1.** Stiffened Moduli by Distance Significantly Change by MVP Status

| | Stiffened Moduli | | |
|---|---|---|---|
| | Normal | MVP | |
| Distance from Cell Boundary | Mean ± SD (Pa) | Mean ± SD (Pa) | Tukey $p$ Value* |
| 0–10 $\mu$m | $2.249 \times 10^5 \pm 7.940 \times 10^6$ | $1.500 \times 10^5 \pm 5.158 \times 10^6$ | 1.000 |
| 10–20 $\mu$m | $7.401 \times 10^4 \pm 3.345 \times 10^6$ | $7.773 \times 10^2 \pm 6.343 \times 10^3$ | 1.000 |
| 20–30 $\mu$m | $1.962 \times 10^6 \pm 5.929 \times 10^7$ | $7.244 \times 10^2 \pm 8.207 \times 10^3$ | $< 0.001$ |
| 30–40 $\mu$m | $3.498 \times 10^2 \pm 5.556 \times 10^3$ | $7.657 \times 10^2 \pm 7.824 \times 10^3$ | 1.000 |
| 40–50 $\mu$m | $1.070 \times 10^2 \pm 2.824 \times 10^1$ | $8.943 \times 10^2 \pm 6.645 \times 10^3$ | 1.000 |
| 50–60 $\mu$m | $1.216 \times 10^2 \pm 4.089 \times 10^1$ | $1.362 \times 10^2 \pm 8.675 \times 10^1$ | 1.000 |
| 60–70 $\mu$m | $1.247 \times 10^2 \pm 4.902 \times 10^1$ | $1.409 \times 10^2 \pm 8.944 \times 10^1$ | 1.000 |
| 70–80 $\mu$m | $9.892 \times 10^1 \pm 1.681 \times 10^1$ | $1.715 \times 10^2 \pm 1.627 \times 10^2$ | 1.000 |
| 80–90 $\mu$m | $1.072 \times 10^2 \pm 1.954 \times 10^1$ | $1.322 \times 10^2 \pm 8.880 \times 10^1$ | 1.000 |
| 90–100 $\mu$m | $1.035 \times 10^2 \pm 1.348 \times 10^1$ | $1.015 \times 10^2 \pm 8.321 \times 10^0$ | 1.000 |

*Tukey $p$ values were calculated from the corresponding row comparisons with a two-way ANOVA and Tukey post test.
The two-way ANOVA also found that $p$ = 0.01166 for a difference in MVP status and $p$ = 0.09257 for a difference in distance.

**Table S2.** Degraded Moduli Significantly Changes by MVP Status and Distance from the Cell Boundary

| | Degraded Moduli | | |
|---|---|---|---|
| | Normal | MVP | |
| Distance from Cell Boundary | Mean ± SD (Pa) | Mean ± SD (Pa) | Tukey $p$ Value* |
| 0–10 $\mu$m | 18.56 ± 9.106 | 17.51 ± 10.28 | 0.1589 |
| 10–20 $\mu$m | 21.20 ± 8.656 | 21.68 ± 9.392 | 0.9820 |
| 20–30 $\mu$m | 19.51 ± 9.211 | 22.81 ± 9.094 | $< 0.001$ |
| 30–40 $\mu$m | 18.20 ± 9.373 | 21.17 ± 9.408 | $< 0.001$ |
| 40–50 $\mu$m | 17.24 ± 9.310 | 19.35 ± 10.00 | $< 0.001$ |
| 50–60 $\mu$m | 16.58 ± 9.128 | 19.15 ± 9.595 | $< 0.001$ |
| 60–70 $\mu$m | 16.07 ± 9.084 | 19.36 ± 8.819 | $< 0.001$ |
| 70–80 $\mu$m | 16.03 ± 8.937 | 23.79 ± 10.16 | $< 0.001$ |
| 80–90 $\mu$m | 15.94 ± 8.721 | 34.13 ± 0.6185 | 0.3416 |

*Tukey $p$ values were calculated from the corresponding row comparisons with a two-way ANOVA and Tukey post test.
The two-way ANOVA also found that $p < 0.001$ for a difference in MVP status and $p < 0.001$ for a difference in distance.

**Table S3.** Hydrogel Strain Energy Density Significantly Changes by MVP Status and Distance from the Cell Boundary

| | Strain Energy Density | | |
|---|---|---|---|
| | Normal | MVP | |
| Distance from Cell Boundary | Mean ± SD (Pa) | Mean ± SD (Pa) | Tukey $p$ Value* |
| 0–10 $\mu$m | 1.103 ± 2.034 | 1.364 ± 1.912 | $< 0.001$ |
| 10–20 $\mu$m | 0.5511 ± 1.324 | 0.4295 ± 0.9011 | $< 0.001$ |
| 20–30 $\mu$m | 0.1706 ± 0.5935 | 0.1387 ± 0.3611 | $< 0.001$ |
| 30–40 $\mu$m | 0.05540 ± 0.2991 | 0.05419 ± 0.1494 | 1.000 |
| 40–50 $\mu$m | 0.02069 ± 0.08755 | 0.02113 ± 0.09472 | 1.000 |
| 50–60 $\mu$m | 0.01219 ± 0.02726 | 0.01079 ± 0.02052 | 0.9987 |
| 60–70 $\mu$m | 0.01027 ± 0.01987 | 0.01024 ± 0.01843 | 1.000 |
| 70–80 $\mu$m | 0.009558 ± 0.01681 | 0.01024 ± 0.01843 | 1.000 |
| 80–90 $\mu$m | 0.01082 ± 0.02022 | 0.01619 ± 0.03974 | 0.7366 |
| 90–100 $\mu$m | 0.01564 ± 0.07038 | 0.01853 ± 0.06382 | 1.000 |

*Tukey $p$ values were calculated from the corresponding row comparisons with a two-way ANOVA and Tukey post test.
The two-way ANOVA also found that $p < 0.001$ for a difference in MVP status and $p < 0.001$ for a difference in distance.
